# Sperm gatekeeping: 3D imaging reveals a constricted entrance to zebra finch sperm storage tubules

**DOI:** 10.1101/451484

**Authors:** T. Mendonca, A. J. Cadby, N. Hemmings

## Abstract

Females across many internally fertilising taxa store sperm, often in specialised storage organs in their reproductive tracts. In birds, several hundred sperm storage tubules exist in the utero-vaginal junction of the oviduct and there is growing evidence that sperm storage in these tubules is selective. The mechanisms underlying female sperm storage in birds remain unknown due to our limited ability to make three dimensional, live observations inside the large, muscular avian oviduct. Here, we describe a new application of fluorescence selective plane illumination microscopy to optically section oviduct tissue from zebra finch *Taeniopygia guttata* females label-free, by harnessing tissue autofluorescence. Our data provide the first description of the 3D structure of sperm storage organs in any vertebrate and reveal the presence of gate-like constricted openings that may play a role in sperm selection.

**Statement of Significance:** Female birds can store sperm in microscopic tubular structures in their reproductive tract for up to several months, depending on species. Studying these sperm storage tubules has been a major challenge due to the muscular and opaque nature of reproductive tracts in birds. We have developed a new method for imaging live reproductive tract tissue using selective plane illumination microscopy, a fluorescence microscopy technique. From these images, we could extract three-dimensional measurements of sperm storage tubules and found these structures to have a gate-like constriction, providing evidence that females can actively select sperm at storage and ultimately influence the paternity of her offspring. Understanding these reproductive adaptations can help improve captive breeding programs and similar conservation strategies.

## Introduction

Across many internal fertilisers, females have evolved the capacity to maintain viable sperm in specialised sperm storage organs in their reproductive tract as a strategy to maximise fertility. Sperm storage ensures the female has sufficient sperm for fertilisation when copulation and ovulation are not synchronised (1). Since female promiscuity is common across taxa (e.g. birds (2), mammals (3–5), reptiles (6), fishes (7) and insects (8–10)), storage also provides the opportunity for females to exert control over post-copulatory processes (11–13). Post-copulatory sexual selection has driven the diversification of sperm storage organs, which vary from single bean-shaped structures in damselflies (14, 15) or one or more sac-like spermathecae in certain fly species (16) (17), to multiple epithelial crypts in snakes (18), lizards (19), turtles (20) and birds (21, 22).

In birds, epithelial sperm storage crypts are called sperm storage tubules (SSTs) and are located in the utero-vaginal junction (UVJ) of the oviduct (23). The number of SSTs possessed by a single female ranges from around 500 in the budgerigar (*Melopsittam undulates*) to 20,000 in the turkey (*Meleagris gallopavo*) (24). A growing body of evidence suggests that avian SSTs are an important site of sperm selection. Steele and Wishart (25) experimentally demonstrated that chicken (*Gallus domesticus*) sperm without surface membrane proteins could not enter the SSTs after normal intra-vaginal artificial insemination, even though these sperm were capable of fertilising the ovum when inseminated beyond the vagina and UVJ. Bobr *et al*. (23) also noted a lack of abnormal sperm in chicken SSTs, suggesting that abnormal sperm are unable to reach or enter sperm storage sites. The large number of SSTs present in the avian oviduct may also allow spatio-temporal segregation of sperm from competing ejaculates (26–28).

Despite evidence that SSTs may act as a filter for high quality sperm, the mechanisms by which sperm are selected at the time of storage remain poorly understood, and how sperm enter and exit the SSTs is unknown. Froman (29) proposed a model where sperm motility, rather than SST function, is pivotal in sperm retention in SSTs. According to this model, sperm must maintain an optimum swimming velocity to maintain their position and counter a fluid current within the SST. This model was supported by evidence that faster sperm emerged out of SSTs later than slower sperm (30), and that passive loss of sperm from storage might be sufficient to explain last male precedence in the domestic fowl, turkeys, and zebra finches (*Taeniopygia guttata*) [31,32; but see 27]. However, there have been no published observations of sperm swimming inside the SSTs, and our own unpublished observations suggest sperm are not motile in storage (see Supplementary Material). Several studies have detected the presence of sperm motility suppressors such as lactic acid in Japanese quail (*Coturnix japonica*) SSTs (33), calcium and zinc in the SSTs of chicken, turkeys and Japanese quail (34, 35), and carbonic anhydrase in the SSTs of turkeys, common quail (*Coturnix coturnix*) and ostriches (*Struthio camelus*) (36–38), and the neurotransmitter acetylcholine, released by nerve endings detected in the vicinity of SSTs (39), has been shown to enhance sperm motility (40), implying a nervous control on sperm mobilisation at ejection from SSTs. Additionally, Hiyama *et al*. (41) presented evidence for the potential role of heat shock protein 70 (HSP70) (42) in enhancing sperm motility at the point of sperm release. The presence of such sperm motility suppressors and activators within or near the SSTs suggests that release of sperm from storage may not be as passive as Froman (29) suggested.

Rather than acting as passive refugia, SSTs may instead be dynamic structures, capable of active constriction and dilation to mediate the entrance and exit of sperm. Although numerous studies have failed to find smooth muscle fibres or myoepithelial cells (39, 43, 44) around SSTs, Freedman *et al*. (39) detected fibroblast-like cells and an F-actin rich cytoskeletal mesh called the “terminal web” in turkey SST epithelia. The terminal web is composed of contractile proteins (actin and myosin) and has been shown to contribute to contractility in other tissues, such as intestinal brush border cells (45, 46) and embryonic pigmented epithelia in chickens (47). Freedman *et al*. (39) also found terminal innervations in the turkey UVJ, suggesting there may be some degree of nervous control over SST function. Recent evidence also suggests the possibility of SST contraction, influenced hormonally by the action of progesterone (48, 49). It is therefore possible that the passage of sperm into and out of storage is controlled, to some degree, by the physical structure of SSTs themselves.

Our understanding of how SST structure influences sperm storage is limited by our relatively basic knowledge of SST morphology. The avian oviduct is convoluted, with opaque, muscular walls, creating numerous practical limitations for making observations of tubules in living epithelial tissue using conventional microscopy techniques. Empirical studies of SST morphology have so far used histology (23, 50, 51) and electron microscopy (34, 43, 52) on fixed tissue sections, but these approaches not only remove functional information, but typically provide two-dimensional information only. Moreover, serial sectioning is laborious, and loss of material can be difficult to avoid. Commonly used light microscopy techniques rely on thin sections and squash preparations (26, 53), which are inappropriate for large tissue samples since they distort structures of interest and allow only limited imaging depths.

In this study we developed a novel method for live, *ex vivo* 3D imaging of SST structure using selective plane illumination microscopy (SPIM). SPIM is highly suitable for imaging large samples at cellular resolution and has lower phototoxicity levels than with other optical sectioning methods which makes it a viable option for imaging living tissue (54, 55). Using SPIM, we were able to optically section UVJ mucosal tissue up to depths of 100 µm without distorting or damaging their structure. We provide the first quantitative estimates of the 3D structure of sperm storage tubules in living tissue, including the relationship between SST length and diameter, and report the existence of a previously undescribed gate-like constriction at the entrance to tubules that may act to regulate sperm transport into and out of storage.

## Materials and Methods

### Animals

This study was approved by the University of Sheffield, UK. All procedures performed conform to the legal requirements for animal research in the UK and were conducted under a project licence (PPL 40/3481) issued by the Home Office.

Zebra finches were from a captive population kept at the University of Sheffield (56, 57). Females (all between one to three years old) were placed in unisex housing for at least two weeks before being paired with males, in double-cages (dimensions of each individual cage: 0.6m × 0.5m × 0.4m) separated by a wire-mesh with the male and female on either side. Each double-cage had a modified nest-box, also with a wire-mesh partition, to allow both birds to enter the nest. This set up allowed the male and female to establish a normal breeding pair bond and enter breeding condition, while preventing them from copulating and therefore ensuring the female had no sperm in her SSTs (sperm can be stored for up to 12 days after mating in zebra finches (24)). Females were only included in the study once they had started to lay eggs, to ensure their oviduct was in full reproductive condition. After they laid their second egg, females were euthanized (in accordance with Schedule 1 [Animals (Scientific Procedures) Act 1986)] and dissected immediately.

### Sample preparation

The oviduct, including the cloaca, was immediately removed from the female and the connective tissue surrounding it was cleared to uncoil and straighten the vagina and the UVJ. The lower end of the oviduct was cut through the middle of the uterus to obtain a segment that included the UVJ, vagina and cloaca. This piece of the oviduct was then cut open lengthwise and pinned flat on a petri-dish filled with silicone elastomer (SYLGARD^®^ 184; Dow Corning). A sufficient quantity of Ham’s F10 Nutrient Mix (Invitrogen, UK) was added to keep the tissue moist but not submerged. For SPIM imaging, UVJ folds were cut individually with iris scissors and mounted one at a time, on a custom-made sample holder (see Supplementary Material) using fine insect needles. The sample holder, with the UVJ fold mounted, was immersed in phenol-free DMEM/F12 media, at 37° C during imaging.

### SPIM imaging

Live UVJ tissue samples, prepared as above, were imaged using a custom-built SPIM microscope (at the University of Sheffield) with laser excitation at 473 nm and a 520 nm long pass (LP) fluorescence emission filter (Semrock, Inc.). The microscope hardware and optical components was based on the OpenSPIM platform (58) but with modifications detailed in Mendonca *et al*. [59; supplementary materials]. The camera, detection and illumination objectives, and magnification were fixed for the system, ensuring that the imaging results were reproduceable. The autofluorescence image stacks were acquired using 500 ms exposure, starting at the outer surface and moving up to 100 µm deep into the tissue fold.

### Characterisation of autofluorescence

SSTs were clearly identifiable in live UVJ tissue during fluorescence imaging on the SPIM and had a punctate appearance on account of autofluorescent granules (Figure 1) which appeared to be mostly confined to SST epithelial cells and were present along the entire length of the SST from orifice to blind end. No other cell structure or organelle was visible in these autofluorescence images.

**Figure 1:**
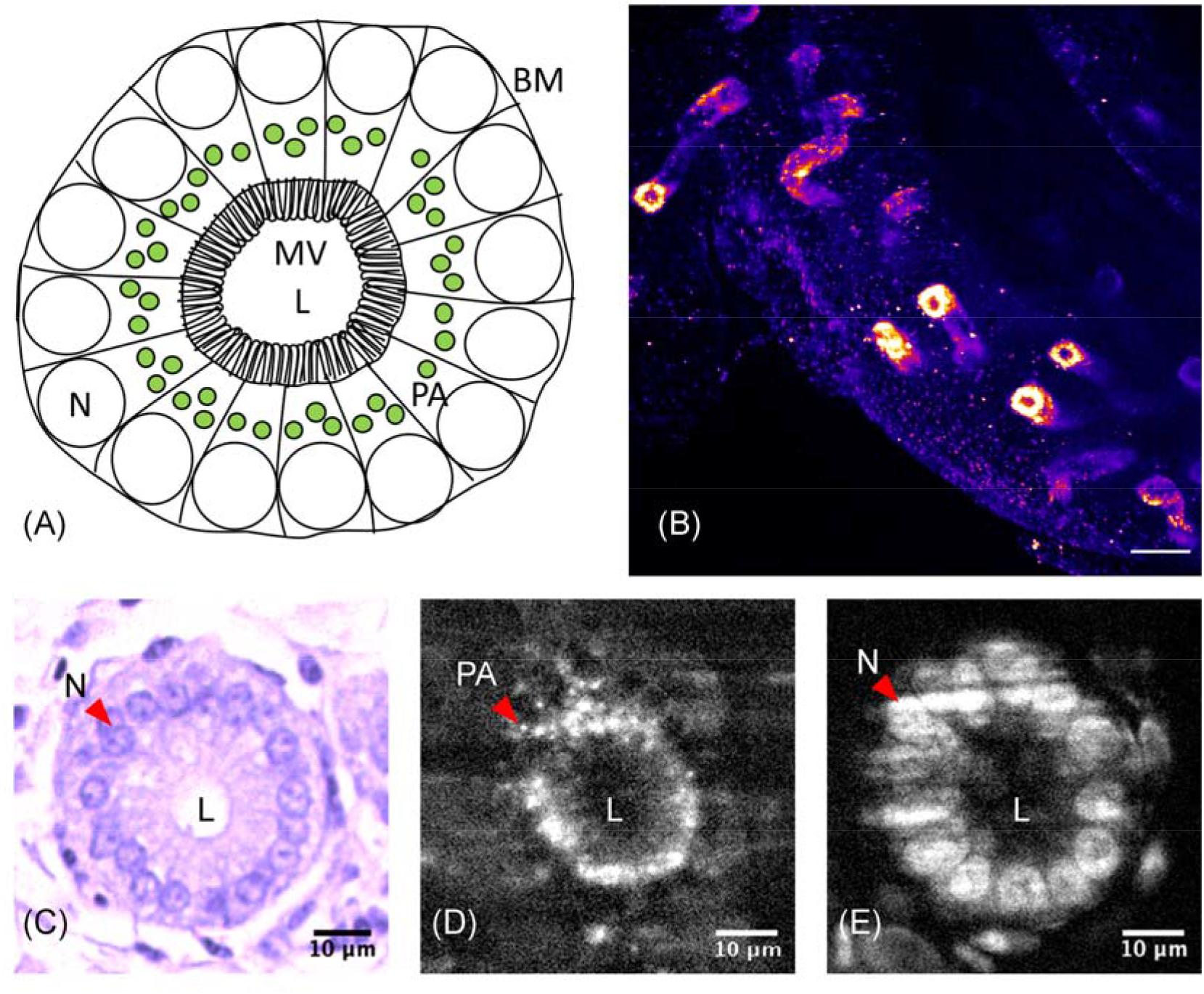
**(A)** Schematic of SST transverse section showing cellular polarisation with nuclei (N) towards the basement membrane (BM) and microvilli (MV) at the apical end of the epithelium. The punctate autofluorescence (PA) detected by the SPIM is present proximal to the nucleus but not at the apical end of the epithelium near the lumen (L). **(B)** Max intensity projection of a UVJ fold with multiple detectable SSTs imaged on the SPIM. Scale – 50 µm **(C)-(E)** Cross section of SST from histology, autofluorescence imaged on the SPIM, and SYTO-13 labelled nuclei imaged on the SPIM respectively. Scale – 10 µm.

To determine the organisation of the autofluorescent granules in the SST epithelium, label-free images from live UVJ folds (n = 10 birds) were compared to those of fixed UVJ folds that had been stained for nucleic acids (n = 3 birds). Folds were also examined on a bright-field microscope after histological sectioning and general histochemical staining (n = 3 birds) (Supplementary Material).

### Image Analysis

The image stacks acquired using the SPIM were used to reconstruct 480 µm × 480 µm × 100 µm tissue sections containing 3D information on SST structure (n = 10 females, one SST per female). SST shape information was extracted by measuring the diameter enclosed by the autofluorescence from images of live tissue, at ten equidistant points along the length of the SST.

UVJ tissue image stacks were first pre-processed in Fiji (60). Individual unbranched SSTs were selected from each female such that the entire SST structure was included in the 3D image stacks. The SSTs follow convoluted paths through the UVJ fold tissue, so in order to measure cross-sectional diameter at multiple points, it was necessary to slice the image volume at arbitrary angles to ensure the measurement planes were perpendicular to the direction of the SST structure. This was accomplished by first tracing the direction of the SST structure using a dilated version of the SST image (generated using the ‘MorphoLibJ’ plugin (61) followed by the application of Gaussian blur), to smooth the punctate autofluorescence. Two outlines for each SST were then semi-automatically traced in 3D, from the orifice to the blind end and along opposite sides of the SST lumen, using the ‘Simple Neurite Tracer’ (62) plugin in Fiji.

The next stage of image analysis was performed using MATLAB^®^ (2015b, version 8.6, MathWorks, Natick, MA). An average trace which passed through the SST lumen was computed from the two traces for each SST. SST lengths were measured from these average traces. For each SST (n = 10), the average trace was interpolated at ten equidistant points (the first at the orifice and the tenth point before the blind end of the tubule) and at each interpolated point a vector describing the direction of the SST at that point was computed using its nearest neighbouring points on the trace (Figure 2). By using these vectors and the interpolated points, slicing planes normal to the vectors were defined. The indices for these slicing planes were used to extract 2D image sections from the un-dilated original image stacks using the ‘ExtractSlice.m’ (63) function. For every extracted slice, its distance from the orifice along the luminal trace of the SST was computed using the ‘Arclength.m’ (64) function, and the major axis diameter (d1) and the minor axis diameter (d2) of the SST (enclosed by autofluorecence) were measured.

**Figure 2:**
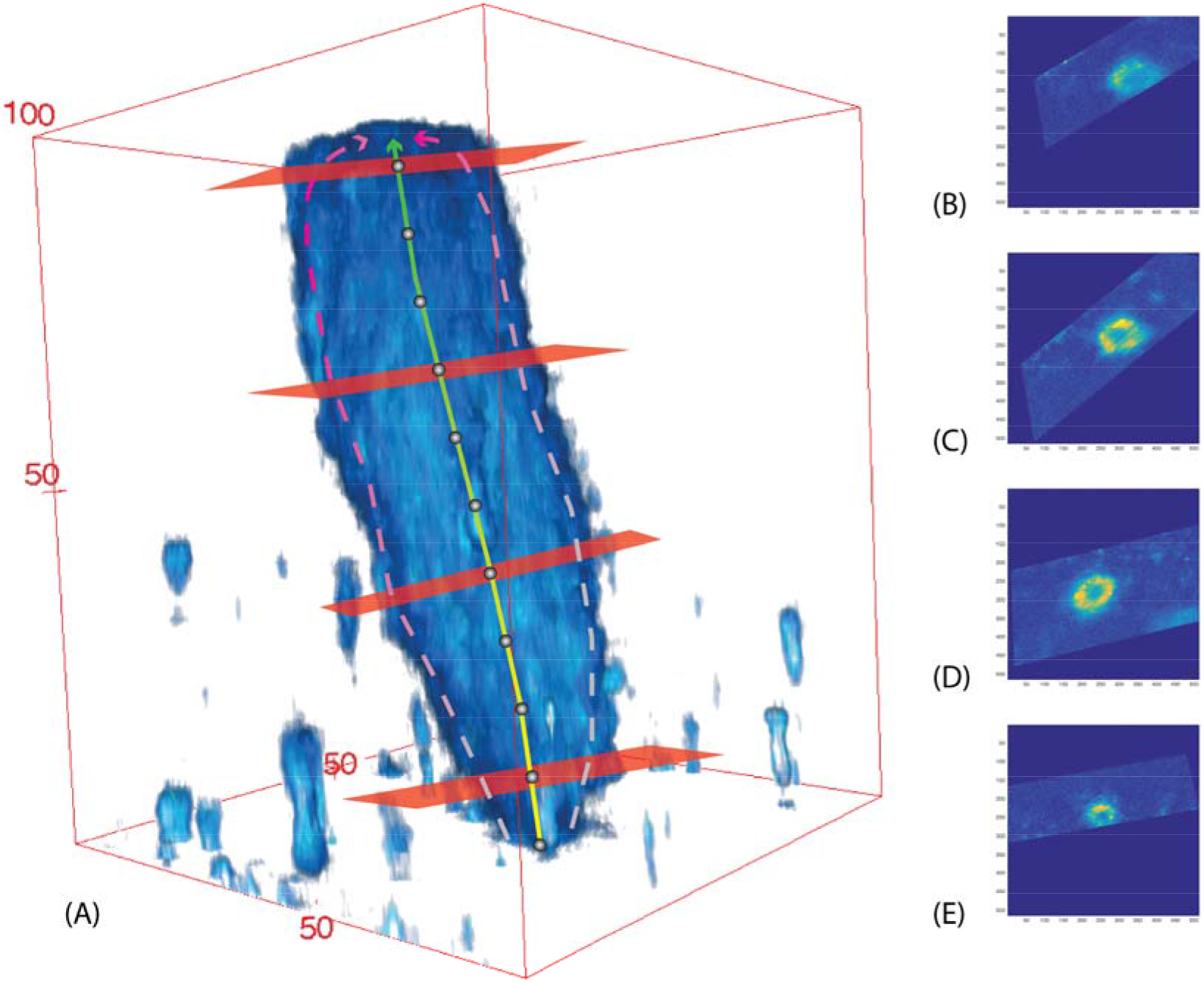
Illustration of the image analysis pipeline. **(A)** 3D rendering of an SST overlaid with traces along the sides (dotted lines), the computed trace through the centre and slices perpendicular to the direction of the SST at 4 example positions (SSTs were sampled at 10 positions represented by grey dots). **(B-E)** Corresponding slices through the SST. Measurements were taken of the major axis and minor axis diameter for each of the slices. Axis units in pixels (converted to microns before data analysis).

### Statistical Analysis

Data analysis was performed using the statistical package R, version 3.2.3 (65). We tested whether SST diameter varied with SST length using a mixed effects model (‘lmer’ function from the ‘lme4’ package (66) along with the ‘lmerTest’ package (67)) with average SST diameter [(d1+ d2) / 2] at the sampled point as the dependent variable, the distance of sampled point from SST orifice and the SST total length as fixed effects, and the bird ID as a random effect to account for repeated measures from each female.

We also assessed if the SST was elliptical or circular in cross-section (the former providing greater epithelial apical surface area for increased contact with sperm) and whether any such ellipticity changed in response to SST length. A circularity index was first calculated by dividing the major axis diameter (d1) by the minor axis diameter (d2), where a circularity index of one indicates a circular SST cross-section. Data were then analysed via a mixed effects model using the ‘lmer’ function (66), with the circularity index as the dependent variable and the total length of the SST as a fixed effect. The sum of the major and minor axis diameters (d1+ d2) was also incorporated as a fixed effect to account for magnitude of change in diameter along each axis, as well as the distance of sampled point from SST orifice, with an interaction term between them. As before, bird ID was included as a random effect to account for repeated measures from each female.

## Results

The diameter of SSTs was found to be notably constricted at their orifice, suggestive of a structural ‘barrier’ for entry and exit (Figure 3 (A)). Beyond this constricted entrance, SSTs were largely tubular in shape, with diameter increasing marginally along the SST’s length until its mid-point, after which diameter decreased again towards the blind end of the SST. This shape can be described by a significant quadratic relationship between SST diameter and the distance from the SST orifice (estimated effect = −16.761, t = −3.085, p = 0.003; Figure 3 (B)). The relationship between SST diameter and distance from the SST orifice was also found using data from labelled tissue (Supplementary Figure S4), confirming that the shape measured from autofluorescence images was not an artefact resulting from the distribution of the autofluorescence granules. Long SSTs were neither wider nor thinner than short SSTs (estimated effect = 0.038, t = 1.058, p = 0.319).

**Figure 3:**
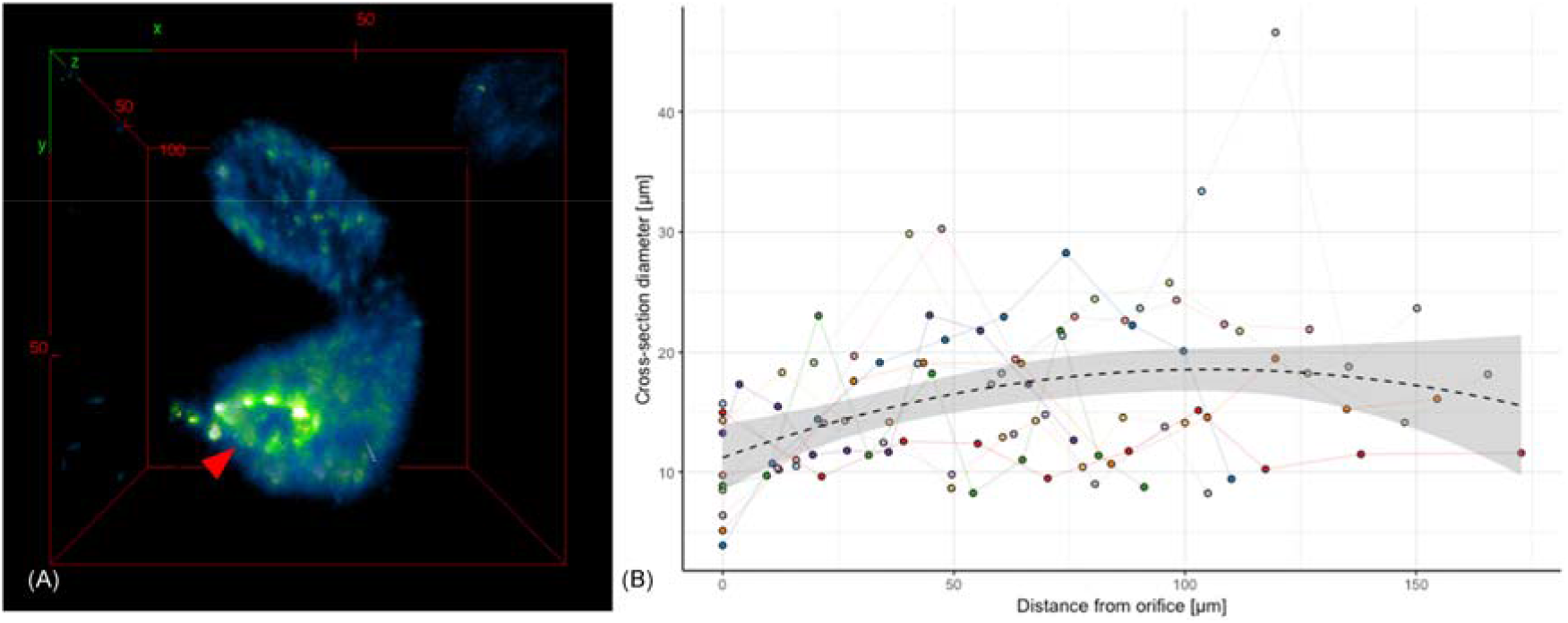
**(A)** 3D rendering of an SST showing its constricted orifice (arrowhead). **(B)** SST diameter has a quadratic relationship with distance from the SST opening suggesting a constriction at the orifice and a slight increase in diameter along its length up to the middle of the SST. Each colour on the plot represents measurements from the same SST (n = 10). Scale on red bounding box is in microns.

SSTs were found to be slightly elliptical in cross-section, with the major axis diameter being 1.642 ± 0.168 times larger than the minor axis diameter. The circularity of the SST in cross-section did not vary significantly with SST diameter (estimated effect = 0.014, t = 0.979, p = 0.330), distance from orifice (estimated effect = 0.003, t = 0.587, p = 0.558), or the interaction between these two variables (estimated effect = −0.00009, t = −0.587, p = 0.559). Circularity was also not related to SST total length (estimated effect = −0.004, t = −1.14, p = 0.282).

Comparisons between the SST measurements from histology and SPIM images indicate that the autofluorescent granules are present in the supranuclear region of the SST epithelium (Figure 1). The size of the lumen diameter scaled linearly with the width of the SST (Supplementary Figure S4), indicating that epithelial cells remained the same thickness in cross-section with increasing SST diameter. This allowed us to extrapolate shape information from the above analyses to the SST lumen, and using this method, we estimated the diameter of the SST orifice to be 3.278 ± 1.115 µm (mean ± SD; Table 1).

**Table 1:** SST dimensions at orifice and widest section. SST diameter is the smallest at its orifice and widest near the middle along its length.

| | Diameter (mean ± SD) | |
| --- | --- | --- |
| | at orifice ($\mu\text{m}$) | at widest section ($\mu\text{m}$) |
| **inter-nuclear diameter<sup>1</sup>** | $12.060 \pm 1.419$ | $30.830 \pm 11.055$ |
| **autofluorescence<sup>1</sup>** | $10.065 \pm 4.288$ | $16.374 \pm 6.610$ |
| **lumen diameter<sup>2</sup>** | $3.278 \pm 1.115$ | $9.128 \pm 1.377$ |

<sup>1</sup>Measurements acquired from SPIM image z-stacks
<sup>2</sup>SST lumen diameter values were predicted from the model describing the relationship between lumen and inter-nuclear diameter.

## Discussion

Using novel 3D imaging methods, we have demonstrated for the first time the existence of a constricted orifice at the entrance/exit of avian SSTs. Such a structure is likely to play an important role in sperm selection at storage. Zaferani *et al*. (68) recently used *in vitro* techniques to demonstrate how constrictions can act as gate-like selective barriers to sperm, allowing only sperm swimming above a threshold velocity to overcome the shear rate at the constriction and pass through. The narrow SST orifices we have found have a mean diameter of ∼3 µm; this, with the added obstruction of microvilli (1-2 µm in length (52)), must act to restrict the rate of sperm (mean diameter at mid-piece - ∼0.6 µm (59)) entering and exiting the SST. We therefore propose that the constricted opening we have found in SST tubules provides a mechanism by which sperm storage and release can be regulated. This supports the idea that avian SSTs play an active and selective role in sperm storage, regulating sperm uptake and release (33–38). The constricted orifice, together with its microvilli, may act as a valve, enforcing unidirectional movement of sperm and preventing them from being flushed back out. The small luminal diameter along the SST (mean: 9 µm, Table 1) may also limit the ability of sperm to turn around inside the SST and swim out.

In terms of overall structure, we found SSTs to be slightly elliptical in cross-section, with the major axis diameter being about 1.6 times larger than the minor axis diameter. This ellipticity was independent of SST radius, the distance along the SST from orifice, or total SST length. Cross-sectional ellipticity increases the surface area of the SST epithelial apical surface, allowing for a greater number of microvilli (as compared to a circular lumen with the same volume) for increased contact with sperm and optimum exchange of nutrients and waste.

We found SST diameter to vary widely across zebra finch SSTs. Birkhead *et al*. (69) suggested that some SSTs might remain inactive in the zebra finch UVJ, even in its fully developed state. It is possible that some of the variation in SST shape that we observed can be explained by the presence of functional and non-functional SSTs, but it is unclear whether thinner, more uniform SSTs, or more distended morphs would represent the active state. Mero and Ogasawara (70), and Burke (71) described ‘swollen’ tubules in chickens and suggested swelling to be associated with sperm release. Such swellings might help explain the outliers in our data (Figure 3 (B)). It is possible that conformational changes in SST shape from functional to non-functional states may be enabled by the F-actin rich terminal web, as seen in turkey SSTs (39), and caused by neural stimulation (39, 72) and/or hormonal effects (48, 49). Variation in SST shape might also be explained by factors not tested in this study, including age, hormone levels and location of the SST in the UVJ.

About 4 – 27% of all the sperm storage tubules in the zebra finch UVJ are branched (28, 73). Branched tubules were not included in our study, but individual branches are expected to show similar shapes as unbranched tubules. Hemmings and Birkhead (28) described sperm from different males differentially stored in separate branches of an SST (albeit a single observation, since in most cases sperm from different males were stored in different SSTs). Further study of the 3D structure of branched SSTs could shed light on mechanisms that prevent sperm mixing in branched tubules.

Our novel 3D data on SST structure were made possible by the presence of punctate/granular autofluorescence, confined to the SST epithelial cells and uniformly distributed throughout the SST’s entire length. These granules were found to have a supranuclear localisation in the SST epithelial cells (Figure 1). Although identifying the exact source of the autofluorescence was beyond the scope of this study, autofluorescence in a similar range has been noted in the ewe (Ovis aries) endometrium (λ_ex_/λ_em_ = 488/525-575 nm) (74) and in human colonic crypts (λ_ex_/λ_em_ = 488/580 nm) (75). While such autofluorescence has been attributed to NADH metabolism in mitochondria (74, 76), another likely source might be lipofuscin in lysosomes (75). Mitochondria are not confined to the apical cytoplasm of SST epithelium as observed in turkeys (77) and chickens (78), so it is unlikely that these granules represent mitochondria. Lysosomes on the other hand, are globular vesicles similar in size (< 1 µm) to the autofluorescent granules observed here (79), and have been detected in the apical cytoplasm of turkey SST epithelia (77), and less abundantly in chickens (78) and passerine alpine accentor (*Prunella collaris*) (79). Multiple studies have also detected the presence of acid phosphatase, an enzyme found in lysosomes, in the supranuclear cytoplasm of SST epithelia in turkeys (21), quail (80), chickens, (81) and ducks (*Anas sp*.) (82), but not in the SST lumen, which corresponds with the autofluorescence pattern we observed here in the zebra finch. Acid phosphatase has been implicated in autolysis associated with oviduct regression (83) as well as with sperm release (82). If this is true, the label-free imaging methods developed here may provide exciting new means for investigating SST functional development throughout the reproductive cycle. Identifying the chemical nature of the autofluorescent substance present in SST granules therefore represents an important avenue for future research.

In summary, we have demonstrated that sperm storage structures in living vertebrate oviductal tissue can be imaged label-free using SPIM microscopy, and this novel 3D imaging technique has enabled us to produce the most detailed account of avian SST structure to date, including the discovery of a previously undescribed gate-like constriction at the entrance/exit of tubules that is likely to act as a key selective barrier. The imaging methods described here hold immense potential for studying *in vivo* sperm storage and sperm-female interactions.

## Authors’ contributions

T.M. coordinated the study, conducted the experiments and analyses, and wrote the manuscript; N.H. conceived the study, conducted the sperm motility experiment and advised on data analyses. All authors participated in the design of the study, helped draft the manuscript and gave final approval for publication.

## Acknowledgements

The authors are grateful to Prof Tim Birkhead and Prof Simon Jones from the University of Sheffield for support. Thanks to Mark Kinch from Skelet.AL, Phil Young, Lynsey Gregory, Emily Glendenning and Jamie Thompson from the University of Sheffield for technical assistance.

