## Supplementary Material 1 for "Sperm gatekeeping: 3D imaging reveals a constricted entrance to zebra finch sperm storage tubules"

**Sperm gatekeeping (Supplementary Material 1: Tissue preparation and comparison of live and fixed tissue)**

T. Mendonca, A. J. Cadby and N. Hemmings

*Sample Mounting*

Conventional sample mounting methods with SPIM involve embedding samples in agarose or alternatively, using adhesives or hooks to grip large tissue samples. Embedding the tissue would block access to SST orifices making it inappropriate for studying sperm-female interactions. Moreover, gripping UVJ tissue with hooks or adhesives was found to be unsuitable in preliminary tests due to the flexible nature of the tissue, which resulted in unwanted tissue movement during imaging scans. To overcome this problem, a modified sample holder was designed for suspending utero-vaginal (UVJ) tissue in the sample chamber as follows:

The bottoms of free-standing micro-centrifuge tubes were cut to a depth of 5 mm. 1 mL syringes were cut to remove the dispensing tip and the micro-centrifuge tube pieces were fixed at right angles to the syringes using epoxy glue (Figure S1). The well of the micro-centrifuge tube was filled with silicon elastomer (SYLGARD^®^ 184; Dow Corning). UVJ tissue samples and/or individual UVJ folds could be mounted on to the surface of the silicon elastomer using the ends of fine insect needles. The free end of the syringe was fastened on the sample arm of the SPIM positioning system for imaging.

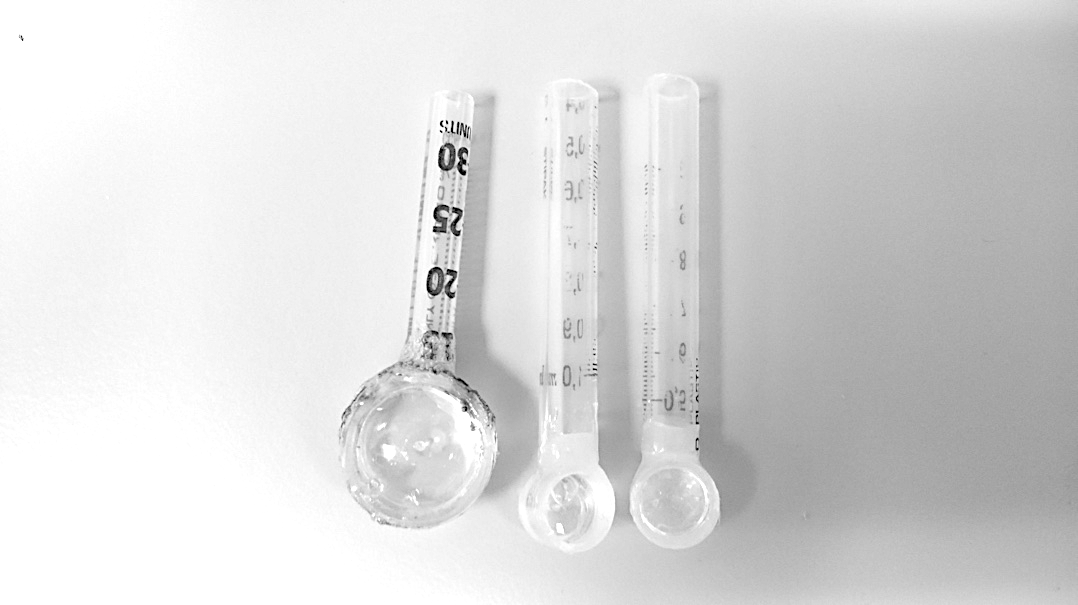

**Figure *S1*:** Customised sample holders of two sizes for imaging oviduct tissue samples. The tissue samples were pinned on the silicon elastomer surface with the UVJ mucosal surface and the SST orifices exposed.

*Localisation of autofluorescence granules*

Live UVJ folds (n = 10 birds) imaged using the SPIM were compared to those of fixed UVJ tissue folds imaged, on the SPIM after staining for nucleic acids (n = 3 birds), and on a bright-field microscope after general histochemical staining (n = 3 birds).

For histochemical examination, pieces of fixed UVJ tissue from three females were sent to the Skeletal Analysis Laboratory (skelet.AL) at the University of Sheffield for processing. The tissue was set in resin blocks and 3 µm thick sections were cut using a microtome. The tissue sections were stained with haematoxylin to label nucleic acids and eosin to label the cytoplasm and other acidophilic structures. The sections were mounted on individual slides and micrographs were taken of the slides on a microscope with bright-field (Leica DMBL with Infinity 3 camera, Luminera Corporation) at 250X magnification using a tiling method. Multiple overlapping image tiles with a 50% overlap were taken for each slide. The individual image tiles were then stitched together using the ‘TrakEM2’ plugin [1] in Fiji [2] to digitally reconstruct each histological section.

10 random SST transverse sections from the histology slides for each bird (n = 3 birds) were measured using Fiji [2]. Two measurements were recorded for each (1) the lumen diameter; (2) the diameter between SST nuclei (referred to as ‘inter-nuclear diameter’ from here on), and (3) the diameter bound by the basement membrane (the ‘outer diameter’) of the SST from each cross-section (Figure S2). The diameter of the SST from autofluorescence images and the inter-nuclear diameter from SPIM images were measured as outlined in the main manuscript text.

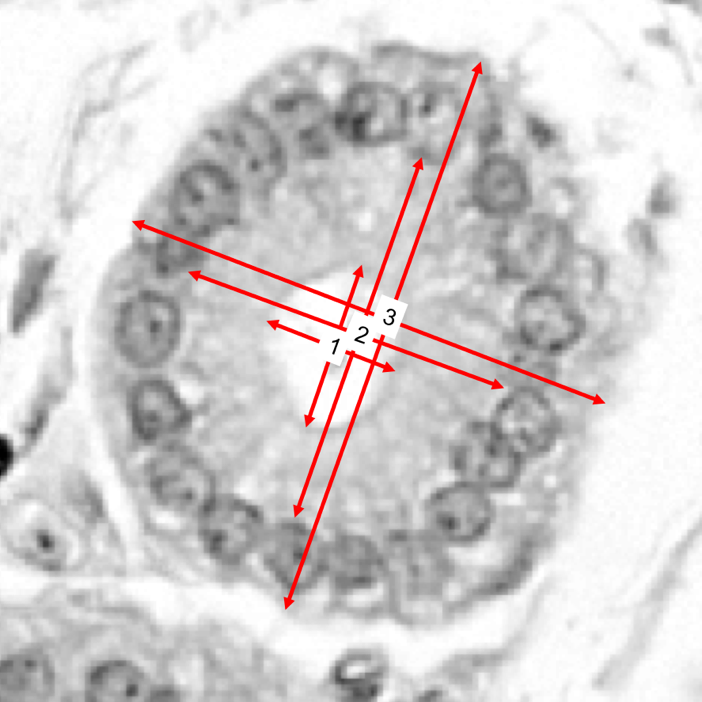

**Figure S2:** Cross section of an SST from histology with arrows indicating the diameters measured from the images: **1**. lumen diameter, **2.** the inter-nuclear diameter and **3.** outer diameter of the SST.

The three different diameter measurements from the histology images showed a linear correlation (lumen and inter-nuclear diameter (log transformed): estimated effect = 1.1339, *t* = 5.479, *p* = <0.0001) suggesting that lumen size scales proportionately with the size of the SST epithelial cells (Figure S3). Changes in the diameter of autofluorescence could therefore be considered to be representative of changes in SST lumen size, so the three-dimensional structure of live SSTs could be analysed from the autofluorescence images.

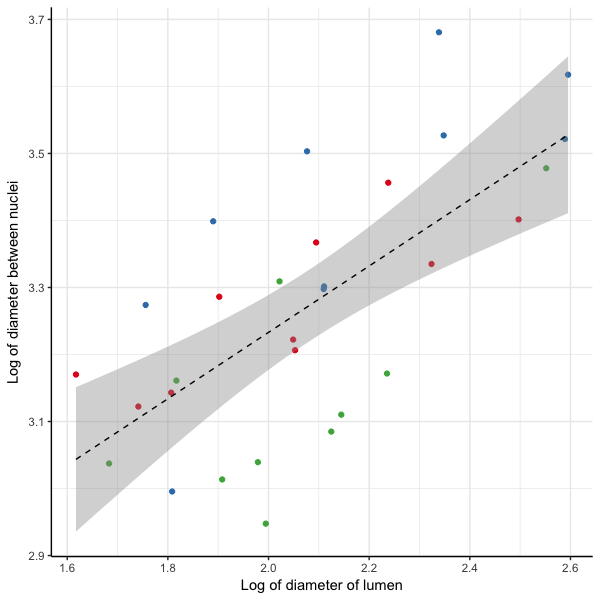

**Figure S3:** Diameter of SST lumen is positively correlated with diameter between SST epithelial nuclei and the outer diameter of the SST. Measurements were recorded from histology images (n = 3 birds, with 10 transverse sections measured per bird). Different colours have been used for data points from each individual.

**Table 1: Diameter measurements [µm] from SST sections**

|  | n  (Birds*) | Diameter  (mean ± S.D.) |
| --- | --- | --- |
| lumen from histology | 3 | 8.299 ± 2.342 µm |
| autofluorescence from SPIM | 10 | 16.075 ± 6.603 µm |
| inter-nuclear diameter from SPIM | 3 | 22.912 ± 7.695 µm |
| inter-nuclear diameter from histology | 3 | 26.873 ± 5.339 µm |
| outer diameter of SST from histology | 3 | 40.235 ± 5.657 µm |

*Data are based on 10 randomly selected SST transverse sections per individual.

*Validation with labelled fixed tissue*

After 2-3 folds had been removed for live tissue imaging, the remaining UVJ tissue was flooded with 5% formalin while still pinned flat on the silicone elastomer and left to fix for two hours at room temperature. The fixed tissue samples were then transferred to micro-centrifuge tubes with 1 mL of 5% formalin for storage at room temperature. For imaging fluorescently stained SST epithelia, individual folds were cut from the fixed UVJ tissue of three females and incubated with 100 µL solution of 10 µM SYTO 13 nucleic acid stain (Molecular Probes Inc., UK) in PBS overnight at room temperature in the dark. The labelled folds were then mounted on the sample holder, one at a time, in the same way as the live tissue samples, and imaged in phenol free DMEM/F12 on the SPIM using the same settings as with live tissue (above), but with 1 ms exposure time.

Image analysis was performed and major axis (d1) and minor axis (d2) diameter measurements were recorded as described for the live tissue image stacks.

A quadratic mixed effects model with average SST diameter [(d1+ d2) / 2] at the sampled point as the dependent variable, the distance of sampled point from SST orifice and the SST total length as fixed effects, and the bird ID as a random effect showed a similar relationship as with measurements from live tissue, between inter-nuclear diameter and distance from SST orifice (estimated effect = -25.293, t = -4.530, p = 0.0001, Figure 12) confirming that the autofluorescence has a uniform distribution and is a fair proxy for analysing SST shape in label-free images (Figure S4).

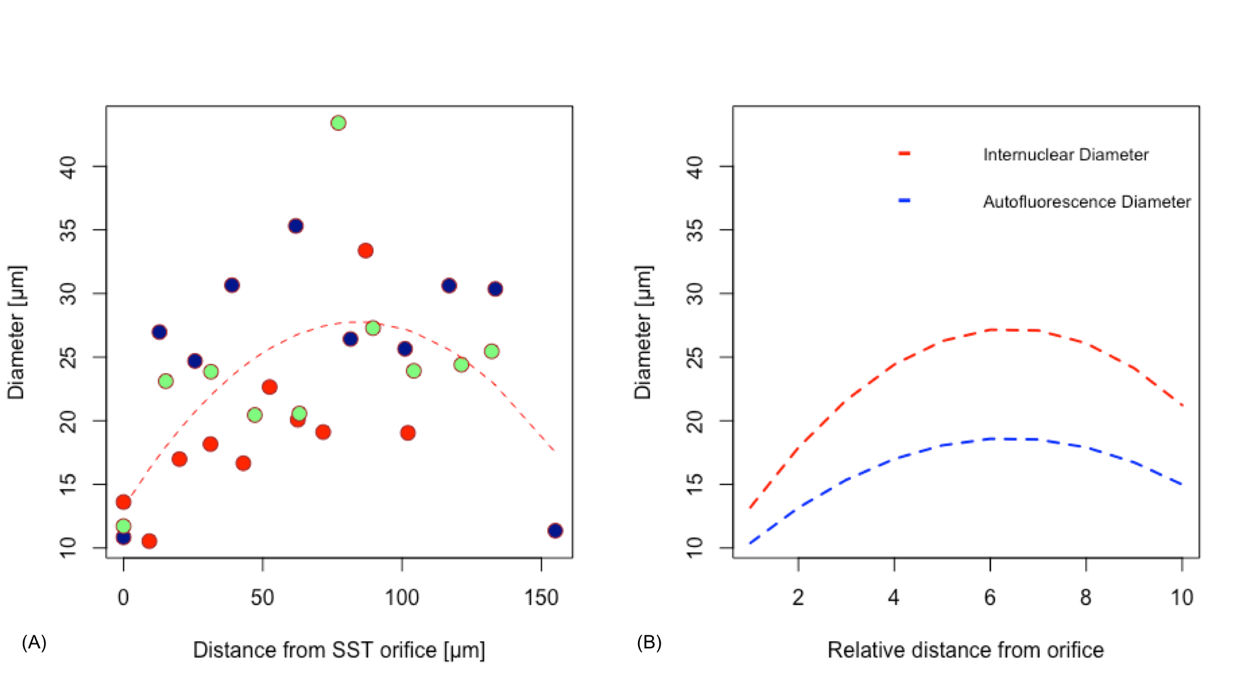

**Figure S4:** The inter-nuclear diameter shows the same relationship with distance from orifice as the autofluorescence diameter. **(A)** Each colour represents data from a SST. **(B)** Fits to experimental data from label-free images of SSTs (blue, n = 10) and labelled fixed SSTs (red, n = 3).
