## Supplementary Material 2 for "Sperm gatekeeping: 3D imaging reveals a constricted entrance to zebra finch sperm storage tubules"

**Sperm gatekeeping (Supplementary Material 2: Sperm motility inside, and after release from, the sperm storage tubules of zebra finches)**

T. Mendonca, A. J Cadby and N. Hemmings

As part of a separate study (by NH), 13 female zebra finches that had copulated with males were dissected to isolate sperm storage tubules (SSTs) for sperm counts using standard light microscopy. During this work, we took the opportunity to observe whether sperm appeared to be motile, both inside and outside the SST.

The females were dissected as described in the main text of the paper, but instead of mounting utero-vaginal folds for SPIM imaging, folds were dissected on a microscope slide under a Nikon SMZ25 stereomicroscope to isolate and open individual SSTs and release sperm following methods described in Hemmings & Birkhead (2017). A single SST containing a large number of sperm (mean = 33 per SST) was isolated per fold (46 SSTs in total), submerged on a microscope slide in warmed nutrient media (Ham’s F10: Invitrogen, UK; typically used for avian sperm motility analysis), which was then observed on a heated stage (38º C) using 200X phase contrast microscopy. Each SST was observed for 5 mins to screen for sperm motility inside the tubule. After this initial observation period, the SST was broken open using fine dissection needles and sperm were released. The slide was then observed for another 5 mins on the heated stage to screen for sperm motility after release from the tubule.

Across at total of 1520 sperm from the SSTs of 13 different females, we did not observe any motile sperm either inside, or after release from, the SSTs. This suggests that sperm are not motile during storage and require some form of activation to trigger motility on release. However, we cannot rule out the possibility that the techniques used for isolating SSTs and extracting sperm may have affected sperm motility, and since the study was not originally designed to test this idea, we do not have data on or observations of stored sperm using other techniques and/or at other places in the oviduct. Our observations should therefore be treated as preliminary and interpreted with care.
